# A hybrid versatile method for state estimation and feature extraction from the trajectory of animal behavior

**DOI:** 10.1101/198879

**Authors:** Shuhei J. Yamazaki, Kazuya Ohara, Kentaro Ito, Nobuo Kokubun, Takuma Kitanishi, Daisuke Takaichi, Yasufumi Yamada, Yosuke Ikejiri, Fumie Hiramatsu, Kosuke Fujita, Yuki Tanimoto, Akiko Yamazoe-Umemoto, Koichi Hashimoto, Katsufumi Sato, Ken Yoda, Akinori Takahashi, Yuki Ishikawa, Azusa Kamikouchi, Shizuko Hiryu, Takuya Maekawa, Koutarou D. Kimura

**Affiliations:** Graduate School of Science, Osaka University, Toyonaka, Osaka 560-0043, Japan; Graduate School of Natural Sciences, Nagoya City University, Nagoya, Aichi, 467-8501, Japan; Graduate School of Information Science and Technology, Osaka University, Suita, Osaka 565-0871, Japan; Department of Polar Science, SOKENDAI, Tachikawa, Tokyo 190-8518, Japan; National Institute of Polar Research, Tachikawa, Tokyo 190-8518, Japan; Department of Physiology, Osaka City University Graduate School of Medicine, Osaka 545-8585, Japan; Center for Brain Science, Osaka City University Graduate School of Medicine, Osaka 545-8585, Japan; PRESTO, Japan Science and Technology Agency (JST), Kawaguchi, Saitama 332-0012, Japan.; Graduate School of Science, Nagoya University, Chikusa, Nagoya, Aichi, 464-8602, Japan; Faculty of Life and Medical Sciences, Doshisha University, Kyotanabe, Kyoto 6100321, Japan; Graduate School of Information Sciences, Tohoku University, Sendai, Miyagi 980-8579, Japan; Atmosphere and Ocean Research Institute, The University of Tokyo, 5-1-5 Kashiwanoha, Kashiwa, Chiba 277-8564, Japan; Graduate School of Environmental Studies, Nagoya University, Nagoya, Aichi 464-8601, Japan; Present address: Department of Ophthalmology, Graduate School of Medicine, Tohoku University, Sendai, Miyagi 980-8574, Japan; Present address: RIKEN Center for Brain Science, Wako, Saitama 351-0198, Japan

## Abstract

Animal behavior is the final and integrated output of the brain activity. Thus, recording and analyzing behavior is critical to understand the underlying brain function. While recording animal behavior has become easier than ever with the development of compact and inexpensive devices, detailed behavioral data analysis requires sufficient previous knowledge and/or high content data such as video images of animal postures, which makes it difficult for most of the animal behavioral data to be efficiently analyzed to understand brain function. Here, we report a versatile method using a hybrid supervised/unsupervised machine learning approach to efficiently estimate behavioral states and to extract important behavioral features only from low-content animal trajectory data. As proof of principle experiments, we analyzed trajectory data of worms, fruit flies, rats, and bats in the laboratories, and penguins and flying seabirds in the wild, which were recorded with various methods and span a wide range of spatiotemporal scales—from mm to 1000 km in space and from sub-seconds to days in time. We estimated several states during behavior and comprehensively extracted characteristic features from a behavioral state and/or a specific experimental condition. Physiological and genetic experiments in worms revealed that the extracted behavioral features reflected specific neural or gene activities. Thus, our method provides a versatile and unbiased way to extract behavioral features from simple trajectory data to understand brain function.

## INTRODUCTION

The brain receives, integrates, and processes a range of ever-changing environmental information to produce relevant behavioral outputs. Therefore, understanding salient behavioral features can augment our understanding of important aspects of environmental information as well as of brain activity, which links the environmental information to behavior. Recent technological development of compact and inexpensive cameras and/or global positioning system (GPS) devices has facilitated convenient monitoring and recording of animal behavior (Brown and de Bivort, 2018; Dell et al., 2014; Egnor and Branson, 2016). However, the behavioral data generated through these approaches are frequently represented as a few simple measures, such as velocity, migratory distance, or the probability of reaching a particular goal, due to the challenges related to identification of specific aspects of behavior to be analyzed; in other words, it is still difficult to figure out how we can describe an animal behavior meaningfully (Berman, 2018). Owing to poor description of behavior, dynamic neural activity, for example, is not sufficiently interpreted even though simultaneous optical monitoring can measure a large number of time-series neural activities (Alivisatos et al., 2012; Landhuis, 2017). This large asymmetry in data richness between neural activity and behavior has emerged as one of the most significant issues in modern neuroscience (Anderson and Perona, 2014; Gomez-Marin et al., 2014; Krakauer et al., 2017).

One way to overcome the challenges in the appropriate descriptions of behavior is to describe its salient features via comprehensive analysis through an approach such as machine learning. Machine learning involves extracting latent patterns and uncovering knowledge from a large amount of data (Bishop, 2006). In fact, multiple behavioral analysis methods based on machine learning have been reported in the last decade (Baek et al., 2002; Branson et al., 2009; Brown et al., 2013; Dankert et al., 2009; Kabra et al., 2013; Mathis et al., 2018; Robie et al., 2017; Stephens et al., 2008; Vogelstein et al., 2014; Wiltschko et al., 2015). Most of these studies have classified behavioral states based on detailed analyses of animal postures as observed in video images (Dell et al., 2014); the classification of behavioral states into classes, such as foraging, sleeping, chasing, or fighting, is considered to be critical for efficient behavioral analysis, as each of behavioral features varies differently across different behavioral states (Egnor and Branson, 2016; Jonsen et al., 2013; Patterson et al., 2008). Although these methods have worked successfully for the analysis of behavioral videos of worms, fruit flies, and rodents in laboratories, they have some limitations. First, these methods are not suitable for analyzing relatively long-distance navigation given their requirement of recording reasonably large and detailed images of animals in the video frame. Second, the extraction of behavioral features from a state, as opposed to just state classification, is more critical in understanding how environmental information and/or brain activities trigger transitions among states for behavioral response.

To analyze relatively long-distance navigation behavior comprehensively, we developed a method for the estimation of behavioral states and extraction of relevant behavioral features based only on the trajectories of animals. For estimating behavioral states, we used an unsupervised learning method involving the expectation maximization (EM) algorithm (Dempster et al., 1977) because it is difficult for the human eye to classify behavior into distinct states without using posture images. For extracting salient behavioral features, we used information gain, an index used for a supervised learning method (the decision tree analysis) (Quinlan, 1992), and compared the features between two different experimental conditions (e.g., with or without certain stimulus). It is because supervised learning is considered advantageous in the extraction of characteristic behavioral features and comparing them among multiple conditions. We named this hybrid supervised/unsupervised machine learning approach as the *st*ate *e*stimation and *f*eature ex*tr*action (STEFTR) method (Fig. 1).

**Figure 1.**
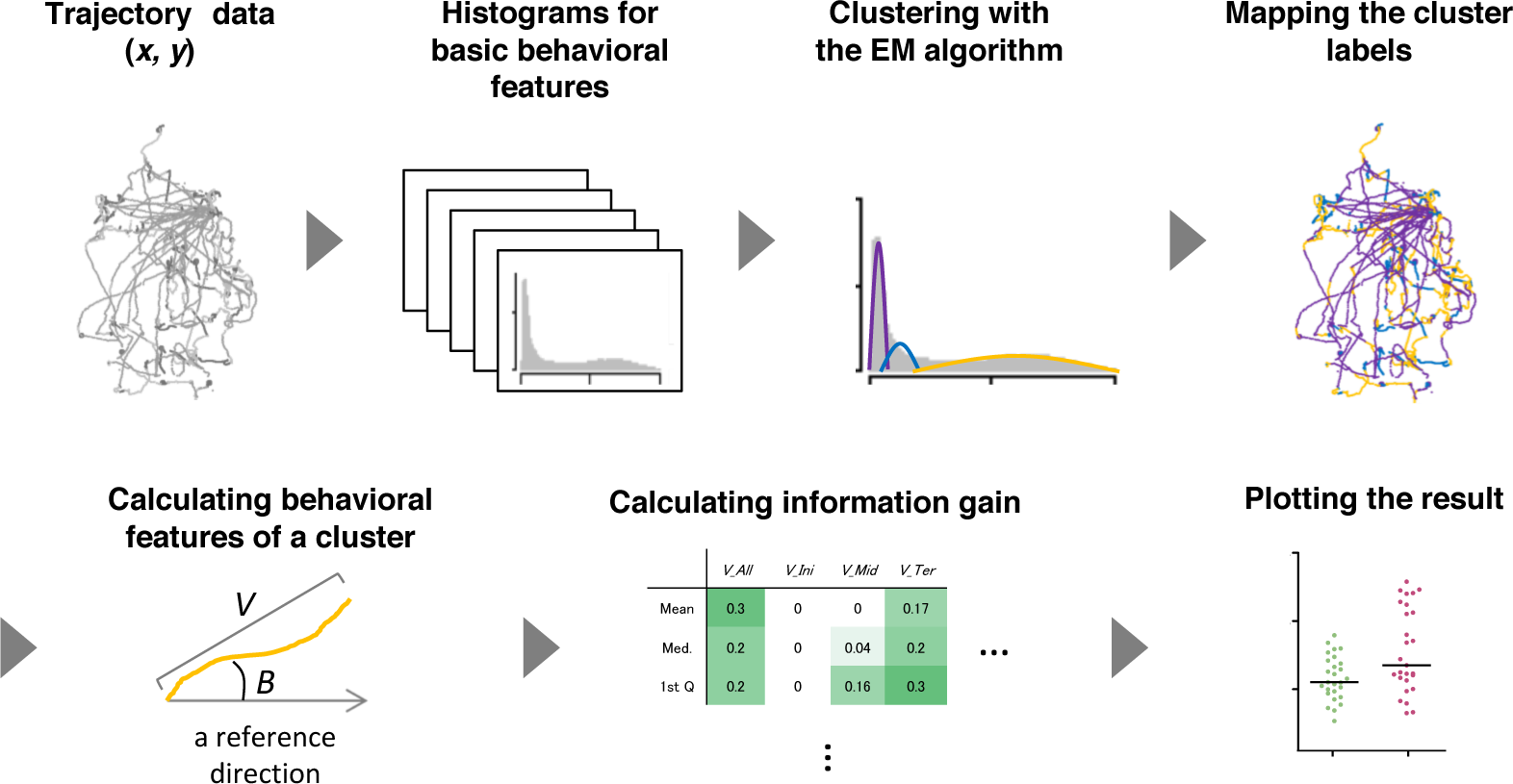
A workflow of the STEFTR method. Trajectory data of animals are used to calculate 8 basic behavioral features, and one of them are analyzed by the EM algorithm to estimate behavioral states (upper panels). From a behavioral state, behavioral features are comprehensively evaluated by using information gain (lower panels).

Because the STEFTR method only uses trajectory information for the analysis, it becomes possible to analyze movement behavior of various animals regardless of the spatiotemporal scale of movement. As proof-of-principle experiments, we analyzed the trajectories of worms, flies, rats, and bats in laboratories and those of penguins and flying seabirds in the wild; these experiments involved a spatiotemporal scale ranging from mm to 1000 km in space and from sub-seconds to days in time. The behavioral states of worms and penguins estimated by the STEFTR method were in reasonable conformation with the ones described in previous literature, supporting the reliability of our method. We further extracted learning-dependent behavioral features from a behavioral state of worms, in which one of the behavioral features is correlated with learning-dependent changes in neural activities. We also analyzed the behavioral features of mutant strains of worms and found that the patterns of features are correlated with gene function, suggesting that comprehensive feature extraction may enable us to estimate unknown functions of a gene product. We were also able to extract learning-dependent features from bats and pheromone-dependent features from fruit flies. Taken together, our findings indicate that the STEFTR method allows us to estimate internal state, neural activity, and gene function related to animal behavior only from movement trajectories, regardless of the recording method or the spatiotemporal scales.

## MATERIAL AND METHODS

### Estimation of behavioral states

For the analysis of trajectory information of an animal obtained from video images or from the GPS device attached to an animal, approximately 1/1,000 and 1/100 of the median recording time across animals were used as an unit for time frame and the time window for moving average, respectively (Table 1). These values were used to draw the 8 histograms of the averages (*Ave*) and the variances (*Var*) of velocity (*V*), bearing (*B*), time-differential of *V* (*dV*) and *B* (*dB*) as the basic behavioral features. The time window for moving average was critical to reduce noise and to detect relatively long trends of behavior. From the histograms, a basic behavioral feature appeared to include multiple normal distributions was selected, and the number of clusters and boundaries of each cluster were automatically determined by EM algorithm (see below). In the case of worms, the cluster analysis was performed with specifying the maximum number cluster of 20. In other cases, maximum cluster number 5 was predetermined based on the knowledge that the number of basic behavioral states are several in general (Patterson et al., 2008).

**Table 1.**
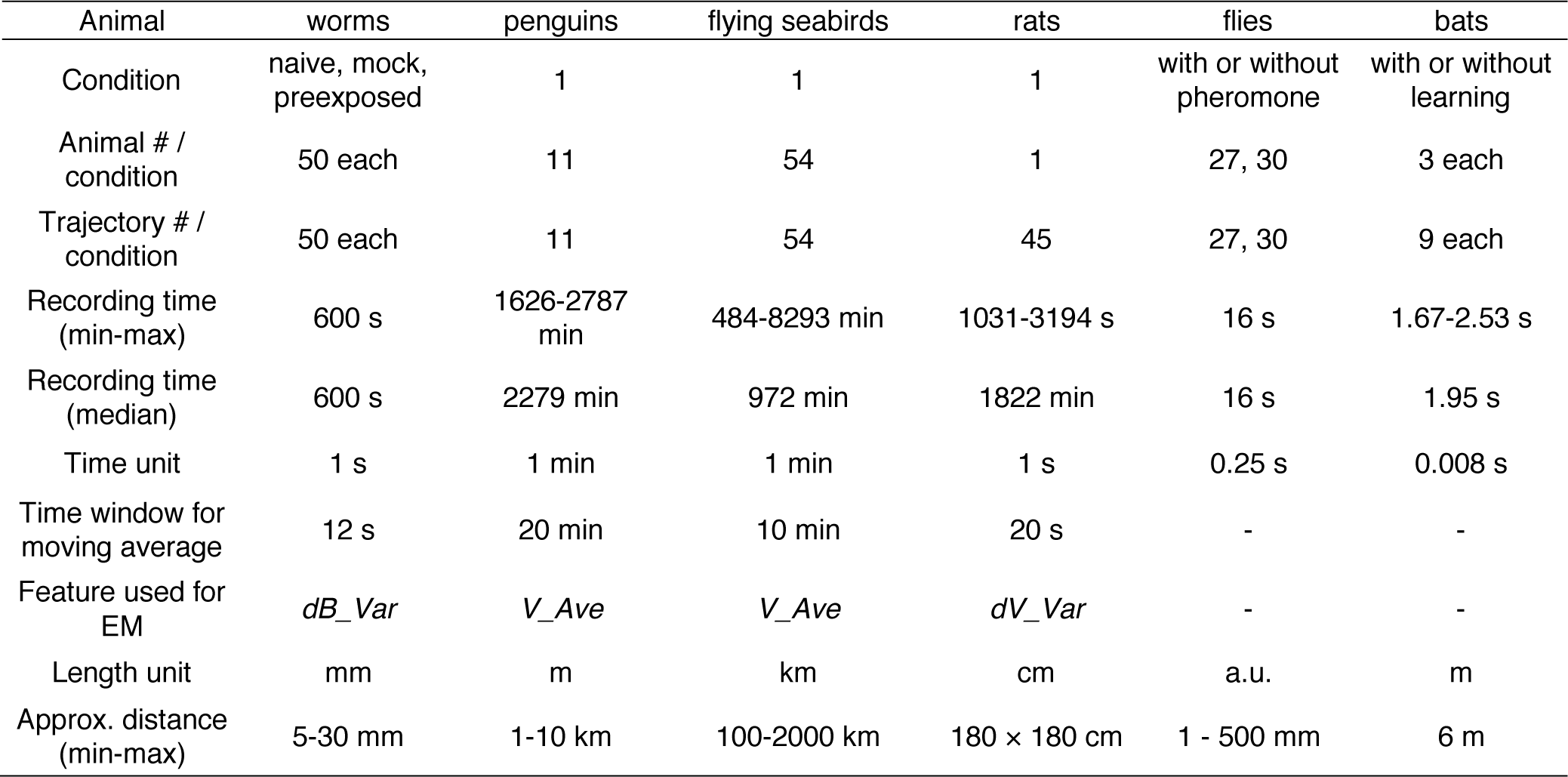
Summary of recording and analysis conditions of animal behavior.

| Animal | worms | penguins | flying seabirds | rats | flies | bats |
| --- | --- | --- | --- | --- | --- | --- |
| Condition | naive, mock, preexposed | 1 | 1 | 1 | with or without pheromone | with or without learning |
| Animal # / condition | 50 each | 11 | 54 | 1 | 27, 30 | 3 each |
| Trajectory # / condition | 50 each | 11 | 54 | 45 | 27, 30 | 9 each |
| Recording time (min-max) | 600 s | 1626-2787 min | 484-8293 min | 1031-3194 s | 16 s | 1.67-2.53 s |
| Recording time (median) | 600 s | 2279 min | 972 min | 1822 min | 16 s | 1.95 s |
| Time unit | 1 s | 1 min | 1 min | 1 s | 0.25 s | 0.008 s |
| Time window for moving average | 12 s | 20 min | 10 min | 20 s | - | - |
| Feature used for EM | *dB_Var* | *V_Ave* | *V_Ave* | *dV_Var* | - | - |
| Length unit | mm | m | km | cm | a.u. | m |
| Approx. distance (min-max) | 5-30 mm | 1-10 km | 100-2000 km | 180 × 180 cm | 1 - 500 mm | 6 m |

The EM algorithm assigns a cluster label to each time frame although the clustering results, which reflect behavioral states of an animal, should be smooth in time. To smooth out the clustering results and removing outlying results, moving average was again applied to the cluster labels, which resulted in clusters resemble to the human-labeled behavioral states.

When the value of a behavioral feature changes suddenly and largely, the influence of the change may extend over a wide range. For example, if an animal moving straightly initiates local search suddenly, *dB* value will be 0°, 0°, 0°, 0°, 0°, 0°, 180°, 0°, 90°, 0°, 270°, etc. If moving average with ±5 time frame was applied, the value change occur from −5 time frame of the sudden value change, which should be compensated. Because worm’s clusters 0 and 1 corresponded to this case, the beginning and the end of each cluster 0 was extended by the half of time window.

The cluster labels obtained as described above were mapped to the corresponding trajectory position with colors. We used a custom-made python program for calculating basic behavioral features, Weka data mining software (the University of Waikato, New Zealand) (Frank et al., 2016) for EM calculation, and Excel (Microsoft) for others.

### EM algorithm for cluster analysis

A set of values of the *i*th basic behavioral feature *F*_*i*_, which were extracted from trajectories of interest, and the number of clusters *N* were given. We employed the EM algorithm to cluster *F*_*i*_ into *N* clusters, i.e., a mixture of *N* Gaussians. The probability distribution of the Gaussian mixtures *M*_*N*_ is represented as follows:

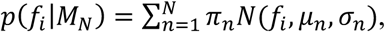

where *f*_*i*_ is a feature value of the *i*th feature, *π*_*n*_ is the mixture weight of the *n*th Gaussian, *μ*_*n*_ is the mean of the *n*th Gaussian, and *σ*_*n*_ is the standard deviation of the *n*th Gaussian. Thus, the EM algorithm was used to estimate the cluster parameters: *π*_*n*_, *μ*_*n*_, and *σ*_*n*_.

### Determination of cluster number using information criteria

The method evaluates a set of clusters (model *M*_*N*_) obtained by the EM algorithm using information criteria, and finds the best *N* using the following criteria.

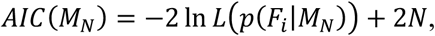

where *L*(*p*(*F*_*i*_|*M*_*N*_) is the maximum likelihood of *F*_*i*_ under model *M*_*N*_ and *N* is the penalty term that measures model complexity.

### Feature extraction with information gain

We leverage information gain to evaluate the classification ability of each feature, i.e., its ability to identify a characteristic of a state (cluster). Information entropy is used to compute the ambiguity of a set of data points according to the following formula:

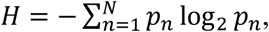

where *p*_*n*_ is the number of data points belonging to the *n*th class (cluster). Given that we classify all the data points into two groups (i.e. two experimental conditions) using a particular threshold related to a specific feature, the feature is considered to be a characteristic feature (in that it classifies the data points well) if the ambiguity within the two groups is lower than that of the original data set. Thus, we evaluated features in terms of their ability in ambiguity reduction upon classification (information gain).

For worm odor avoidance behavior, we extracted behavioral features that have positive information gain in naive versus pre-exposed worms or in mock-treated versus pre-exposed worms. Next, we chose the extracted features that were common for both comparisons; these features were termed as “features modulated in a learning-dependent manner.” For flies, behavioral features were compared between with or without pheromone tapping. For bats, behavioral features were compared between unfamiliar flights (1^st^-3^rd^) and familiar (10^th^-12^th^) flights. The Weka software was used for these calculations.

### Behavioral parameters included in a feature vector

For the machine learning analysis of worm’ odor avoidance behavior, the following behavioral features were calculated for each cluster 0 segment from the coordinates of the centroid of the trajectory: velocity (*V*), bearing (*B*), odor concentration the worm experienced during the run (*C*), the time differential values for these (*dV*, *dB*, and *dC*), directedness (*Dir*) (Gorelik and Gautreau, 2014), curvature (called weathervane; *WV*) (Iino and Yoshida, 2009), and durations of cluster 0 and 1 (*Clst0Dur* and *Clst1Dur*, respectively). For *V*, *dV*, *B*, *dB*, *C*, *dC*, and *Dir*, the average (*Ave*) and median (*Med*) values for at the initiation (*Ini*), middle (*Mid*), termination (*Ter*), and all (*All*) periods of a cluster 0 segment were calculated. A total of 333 features was calculated by combining all these features.

For analyzing changes in the flight of bats, the following behavioral features in each flight were calculated from the coordinates of the bats and obstacles: three-dimensional flight velocity (*V*), horizontal and vertical bearings of the flight (*B_hori* and *B_vert*, respectively), distance (*R_obs*) and bearing (*B_obs*) of the bat to the nearest edge point of the obstacle chain array, longitudinal directional distance to the frontal chain array (*R_x*), and lateral directional distance to the inside pitch of the chain array (*R_y*). Time-differential values were calculated for *V* (*dV*), *B* (*dB*), *dB* (*ddB*), and the flight height (*dH*), which were calculated with frame units of the high-speed video cameras (1/125 s). All flight trajectories were divided into three segments: early, middle, and late terms. The time window for the analysis of each behavioral feature was 0.1 s, 0.2 s, or 0.3 s before or while (*t* = 0) passing through the chain array. A total of 42 features was calculated by combining all these features.

Excel and Visual C# (Microsoft) were used for the calculations, while the Beeswarm package for R (The R Project) was used to obtain a scatter plot of the data.

### Worms

The culture and handling of *Caenorhabditis elegans* strains were performed according to techniques described previously (Brenner, 1974). Wild-type Bristol strain RRID:WB-STRAIN:N2_Male and mutant strains RRID:WB-STRAIN:MT1219 *egl-3(n589)*, RRID:WB-STRAIN:VC671 *egl-3(ok979)*, RRID:WB-STRAIN:CX4544 *ocr-2(ak47),* RRID:WB-STRAIN:JC1636 *osm-9(ky10)*, and RRID:WB-STRAIN:FK127 *tax-4(p678)*, RRID:WB-STRAIN:MT6308 *eat-4(ky5)*, and RRID:WB-STRAIN:IK105 *pkc-1(nj1)* were obtained from the Caenorhabditis Genetics Center at the University of Minnesota, USA. The RRID:WB-STRAIN:KDK1 *dop-3(tm1356)* strain was originally obtained from the National BioResource Project (Japan) and back-crossed with the wild-type N2 strain five times.

A 2-nonanone avoidance assay was performed according to the protocol described previously (Kimura et al., 2010; Yamazoe-Umemoto et al., 2015). Briefly, 2-3 young adult hermaphrodite worms grown synchronously were placed in the center of a 9-cm nematode growth media (NGM) plate. Worm behavior was recorded for 12 min after 2-μL of 30% 2-nonanone (cat. no. 132-04173; Wako, Japan) diluted in 99.5% ethanol (cat. no. 0057-00456; Wako, Japan) were placed at two spots on the surface of the NGM plate. This assay was performed under the following three conditions: (1) Naive—the worms cultivated on 6-cm NGM plates with the RRID:WB-STRAIN:OP-50 bacteria as food were briefly washed with NGM buffer and subjected to the assay; (2) Pre-exposed—the worms were subjected to the assay after being pre-exposed to 0.6 µL of 15% 2-nonanone spotted on the lid of a 6-cm NGM plate for 1 h without food; and (3) Mock—the worms were subjected to the assay after being pre-exposed to ethanol similarly to the pre-exposed condition. We added the mock-treated control group to ensure that the starvation itself did not affect the odor avoidance behavior of worms and to extract behavioral features modulated by odor pre-exposure compared to the naive and mock-treated control groups. Images of worms on the 9-cm NGM plate during the odor avoidance assay were acquired by a high-resolution USB camera (DMK 72AUC02; The Imaging Source, USA) with a lens (LM16JC5MW; Kowa, Japan) at 1 Hz for 12 min. The coordinates of individual animals’ centroids were acquired from the recorded images using the Move-tr/2D software (Library Co., Ltd., Tokyo, Japan) and used for the STEFTR analysis.

Similar to the other sensory behaviors of the worms, trajectories in the 2-nonanone avoidance behavior can be divided into two states: (1) “run”—a relatively long period of straight movement, and (2) “pirouette”—a period of short movements interrupted by frequent reversals and turns (Kimura et al., 2010; Pierce-Shimomura et al., 1999). The angular change per second was calculated from the centroid coordinates, and movements of 1 s with angular changes larger than 90° were classified as a turn. The histogram of turn intervals could be fitted to two exponentials, suggesting that the turn intervals are regulated by two probabilistic mechanisms (Pierce-Shimomura et al., 1999; Yamazoe-Umemoto et al., 2015). The time point at which the two exponentials intersected was defined as *t*_*crit*_, and turn intervals longer or shorter than the *t*_*crit*_ were classified as runs or included in pirouettes, respectively. The *t*_*crit*_ was calculated for the control (i.e., naive and mock-treated) condition for wild-type and mutant strains. Excel (Microsoft) was used for the above calculations. The odor concentrations that the worms experienced at specific spatiotemporal points were calculated according to the dynamic odor gradient model based on the measured odor concentration (Tanimoto et al., 2017; Yamazoe-Umemoto et al., 2018).

Statistical analyses were performed with Prism ver. 5.0 for Mac OSX (GraphPad Software, CA, USA) and R (The R Project). Sample size was determined based on the previous report (Yamazoe-Umemoto et al., 2015). A part of the original data used in this study had already been analyzed and published previously (Yamazoe-Umemoto et al., 2015), and re-analyzed with the STEFTR method.

### Penguins

Fieldwork was performed on chick-rearing Adélie penguins at Hukuro Cove colony (69°13′ S, 39°38′ E) in Lützow-Holm Bay, East Antarctica. GPS-depth loggers (GPL380-DT or GPL400-D3GT, weighing 55-85 g; Little Leonardo, Japan) were deployed among 11 penguins during the period between 27 December 2016 and 10 January 2017 and recovered from all the birds after 1-2 days. While the loggers were set to record GPS positions and depth every second, they could not record GPS positions when the penguins were diving. Therefore, we linearly interpolated the data, when necessary, to obtain GPS positions every 1 minute before further analysis. See Kokubun et al. for methodological details (Kokubun et al., 2015). This fieldwork was carried out in accordance with the recommendations of the Law relating to Protection of the Environment in Antarctica. The protocol was approved by the Ministry of the Environment, Government of Japan. Sample size was not predetermined.

### Flying seabirds

Fieldwork was performed on *Calonectris leucomelas* at Funakoshi-Ohshima Island (39°24’N, 141°59’E) between August and September in 2011, 2012, 2013, and 2015. We attached GPS loggers (GiPSy-2, 37 × 16 × 4 mm or GiPSy-4, 37 × 19 × 6 mm; TechnoSmArt, Roma, Italy) to the back feathers of chick-rearing Streaked Shearwaters with Tesa® tape (Beiersdorf AG; GmbH, Hamburg, Germany) and cyanoacrylate glue (Loctite®401; Henkel Ltd., Hatfield, UK). The loggers were housed in waterproof heat-shrink tubing and set to record one fix per minute. The total weight of the unit was 25 g, which was less than 5% of the mean mass of the birds in accordance with the suggested load limit for flying seabirds. After approximately two weeks of deployment, we recaptured and retrieved the loggers. See Yoda et al. for methodological details (Yoda et al., 2014). The study was carried out in accordance with the recommendations of the guidelines of the Animal Experimental Committee of Nagoya University. The protocol was approved by the Animal Experimental Committee of Nagoya University. Sample size was not predetermined. A part of the original data used in this study had already been analyzed and published previously (Yoda et al., 2014), and re-analyzed with the STEFTR method.

### Rats

Locomotion data of an adult male Long Evans rat were obtained from the Collaborative Research in Computational Neuroscience (CRCNS; RRID:SCR_005608) data sharing website (https://crcns.org, hc-3 dataset, ec013 rat) (Mizuseki et al., 2014). The rat foraged for randomly dispersed water or foods on an elevated open field (180 cm × 180 cm) for 17 - 53 min. The rat’s position was tracked by monitoring two light-emitting diodes mounted above the head with an overhead video camera at 30 Hz. The 30-Hz tracking data were resampled to 39.0625 Hz for offline processing. This study was carried out in accordance with the recommendations of the Regulations on Animal Experiments in Osaka City University. The protocol was approved by the Animal Care and Ethics Committee of Osaka City University. Sample size was not predetermined. The original data used in this study had already been analyzed and published previously (Diba and Buzsáki, 2008; Mizuseki et al., 2009; 2014), and re-analyzed with the STEFTR method.

### Flies

Fruit flies *D. melanogaster* were raised on standard yeast-based media at 25°C and 40 - 60% relative humidity under a 12-hour light/dark cycle. *Canton-S* flies aged between 6-8 days after eclosion were used as a wild-type. After eclosion, the males were housed singly, while females were housed in groups until the experiment.

The locomotion measurement was performed as described previously with minor modifications (Kohatsu and Yamamoto, 2015; Kohatsu et al., 2011). Briefly, a male fly was tethered with a metal wire on its dorsal thorax and positioned over an air-supported Styrofoam ball (diameter, c. a. 6 mm). The locomotion trajectory of the fly was recorded by monitoring the rotations of the Styrofoam ball using an optical computer mouse sensor (BSMRU21BK; BUFFALO INC., Nagoya, Japan). The sensor detected the movements of the ball in the horizontal (Δ*x*) and vertical (Δ*y*) directions, which correspond to lateral and forward movements of the male fly, respectively. The Δ*x* and Δ*y* values, together with timestamps, were sent to a computer at 60 Hz via an Arduino Due microcontroller (Switch Science, Japan) with a custom sketch program. The 60-Hz data were down-sampled to 4-Hz data for the information gain analysis. The measurements were obtained at 25±1°C and 50±10% relative humidity and within 4 hours after light onset.

Female pheromones were applied to the male fly by placing the female’s abdomen in contact with the male’s foreleg at the onset of the measurement. A manipulator (M-3333, Narishige, Tokyo, Japan) actuated a pipette with a volume of 200 µL (FUKAEKASEI Co., Ltd., China), in which a live female with her abdomen exposed toward a male fly was captured. We manually controlled the position of the manipulator to contact the female’s abdomen to the male’s foreleg. This contact procedure was omitted in the control experiments.

Visual stimulus was applied directly after pheromone application by starting horizontal movements of the female fly in front of the male fly as described (Kohatsu et al., 2011). The visual stimulus consisted of ten left-right horizontal movements of the female that lasted for 40 s. Each movement started with the female fly in the front of the male fly (i.e., center) and continued as the female fly moved left until it reached the left end of the rail (i.e., 5 mm away from the center), then moved right until it reached the right end of the rail (i.e., 5 mm away from the center), and ended when it came back to the center with a constant velocity of 5 mm/s. This movement was driven by a stepper motor (42BYG Stepper Motor, Makeblock Co., Ltd., Shenzhen, China) controlled by a custom sketch program (processing software version 3.3.7). We defined one round (4 s in total) as the movement related to the female starting to moving away from the center, reaching the left end of the rail, passing the center, moving away to reach the right end of the rail, and coming back to the center again. Each of the “moving away” and “coming back” periods lasted for 1 s.

The Δ*x* and Δ*y* values with timestamps obtained in the final eight rounds were used for the analysis. To detect the characteristic parameters for the chasing behavior, we used Δ*x* and Δ*y* values during the period when the female was moving away from the male (2 s/round). We set the angle of the chasing behavior as 0 degree when the male moved forward. Angles between 0 and 90 degrees indicate that the male fly is moving towards the female that was moving away from the male.

As the parameters (velocity, bearing, and their time-differential values) were not normally distributed (Shapiro-Wilk test; see Supplementary Table 13), their values were compared between conditions (with/without pheromone) using the Mann-Whitney U test followed by Bonferroni correction for multiple comparison. We used the Steel-Dwass test to compare values of the parameters between rounds. Statistical analyses were conducted using R software version 3.4.4. No statistical methods were used to pre-determine sample sizes, but our sample sizes are similar to those in previous studies (Kohatsu and Yamamoto, 2015; Kohatsu et al., 2011).

### Bats

Three adult Japanese horseshoe bats (*Rhinolophus ferrumequinum nippon*, body length: 6–8 cm, body mass: 20–30 g) were captured from natural caves in the Hyogo and Osaka prefectures in Japan as previously described (Yamada et al., 2016). The bats were housed in a temperature- and humidity-controlled colony room [4 m (L) × 3 m (W) × 2 m (H)] with a 12-h light/dark cycle at Doshisha University in Kyoto, Japan, and were allowed to fly freely and given access to mealworms and water. Captures were conducted under license and in compliance with current Japanese law. This study was carried out in accordance with the recommendations of Principles of Animal Care (publication no. 86-23 [revised 1985)] of the National Institutes of Health) and all Japanese laws. The protocol was approved by the Animal Experiment Committee of Doshisha University.

Methods for acoustic navigation measurement in bats have been described elsewhere (Yamada et al., in revision). Briefly, the experiments were conducted in a flight chamber constructed using steel plates [9 (length) × 4.5 (width) × 2.5 m (height)] under lighting with red filters (>650 nm) to avoid visual effects on the bats. An obstacle environment was constructed using plastic chains (diameter: 4 cm) suspended from the ceiling of the chamber. The chains were arranged at 15-cm intervals along the *x*-axis and at 22-cm intervals along the *y*-axis so that the bat was forced to fly in an S-shaped pattern without passing between chains. Three naive bats were observed for 12 continuous repeated flights so that their echolocation behavior in unfamiliar and familiar spaces could be compared. In this study, the first three flights were defined as unfamiliar flights, while the last three flights were defined as familiar flights.

The flight behavior of the bats was recorded at 125 frames/s using two digital high-speed video cameras (MotionPro X3; IDT Japan, Inc., Japan) placed in the left and right corners of the flight chamber. Based on a direct linear transformation technique, the successive 3D positions of the flying bats, as well as the locations of other objects, were reconstructed using motion analysis software (DIPPMotionPro ver. 2.2.1.0; Ditect Corp., Japan). The statistical calculations were performed with SPSS version 23 (IBM Corp.).

### Calcium imaging of worm’s neurons

Calcium imaging of the worms’ ASH neurons was performed according to the previous method with some modifications (Tanimoto et al., 2017). Briefly, transgenic strains expressing GCaMP3 (Tian et al., 2009) and mCherry (Shaner et al., 2004) in ASH sensory neurons under the *sra-6* promoter (KDK70034 and KDK70072; 20 ng/µl of *sra-6p::GCaMP3*, 20 ng/µl of *sra-6p::mCherry*, 10 ng/µl of *lin-44p::GFP*, 50 ng/µl of PvuII-cut N2 genomic DNA as a carrier in N2 background) were placed on an NGM agar plate on a robotic microscope system, OSB2 (Tanimoto et al., 2017). Although these transgenic worms were immobilized with the acetylcholine receptor agonist levamisole (Lewis et al., 1980) for high-throughput data acquisition through simultaneous imaging of multiple worms, the previous study revealed that the ASH activity is essentially unaffected by levamisole-treatment (Tanimoto et al., 2017). A constant gas flow of 8 mm/min was delivered, in which the mixture rate of 2-nonanone gas with air was changed to create a temporal gradient of odor concentration. The temporal change in odor concentration was measured by a custom-made semiconductor sensor before and after the series of calcium imaging experiments on each day. The fluorescence signals of GCaMP3 and mCherry in ASH neurons were divided into two channels using W-View (Hamamatsu, Japan), an image splitting optic, and captured by an electron multiplying charge-coupled detector (EM-CCD) camera (ImagEM; Hamamatsu, Japan) at 1 Hz. The intensities of fluorescence signals from cell bodies were extracted and quantified by ImageJ (NIH) after background subtraction. The average ratio over 30 s prior to the odor increase was used as a baseline (*F*_*0*_), and the difference from *F*_*0*_ (Δ*F*) was used to calculate the fluorescence intensities of GCaMP3 and mCherry (*F* = Δ*F/F*_*0*_). The ratio between florescence intensities of GCaMP and mCherry (GCaMP/mCherry) was used in the figure.

## RESULTS

### Estimation of behavioral states

As the first part of the analysis, we classified the trajectory into several behavioral states based on the distribution of a basic behavioral feature. Behavior of animals consists of several states (Jonsen et al., 2013; Patterson et al., 2008), where basic behavioral features such as speed and direction change are likely distributed probabilistically with a center value that is optimal for each state. Thus, behavior can be more easily characterized when the behavioral features are analyzed for each state rather than for the entire behavior as a whole. In fact, classifying the trajectory into several states is one of the essential preprocessing steps in trajectory mining of people and vehicles in data science (Zheng, 2015).

For the state classification, we calculated the averages (*Ave*) and variances (*Var*) of four basic behavioral features: velocity (*V*), temporal changes in velocity (i.e. acceleration, *dV*), bearing (*B*), and temporal changes in bearing (*dB*). These 8 features were represented in the form of histograms. Based on the hypothesis that basic behavioral features are likely to be distributed probabilistically for each state, we chose one of the features whose histogram likely consists of multiple probabilistic distributions, and classified the feature values into several clusters with the EM algorithm (Dempster et al., 1977). The EM algorithm, which is an iterative method to estimate model parameters that maximize the likelihood of the model for data, was used to estimate parameters such as the average and variance of each cluster. Each cluster was then regarded as a behavioral state. As proof-of-principle experiments, we analyzed the trajectories of worms and penguins, whose behavioral states have been studied previously using other methods (Pierce-Shimomura et al., 1999; Yoda et al., 2001).

The roundworm *Caenorhabditis elegans* has been used as a model animal for quantitative behavioral analysis owing to the ease of tracking behavior (movement for a few cm on agar surface can be easily recorded with an inexpensive high-resolution camera), optical monitoring neural activities, and genetic analyses and manipulations (De Bono and Maricq, 2005). Further, the neuronal wiring in *C. elegans* has been described in complete detail (White et al., 1986). In this study, we focused on the avoidance behavior to the repulsive odor of 2-nonanone (Fig. 2A, left) (Bargmann et al., 1993; Kimura et al., 2010). We chose this behavioral paradigm for the proof-of-principle experiment because the odor avoidance behavior has been quantitatively, although not fully, analyzed previously (Kimura et al., 2010; Yamazoe-Umemoto et al., 2015). The behavior of the worms was recorded with an USB camera for 12 minutes, and the position of each worm’s centroid was extracted every second.

**Figure 2.**
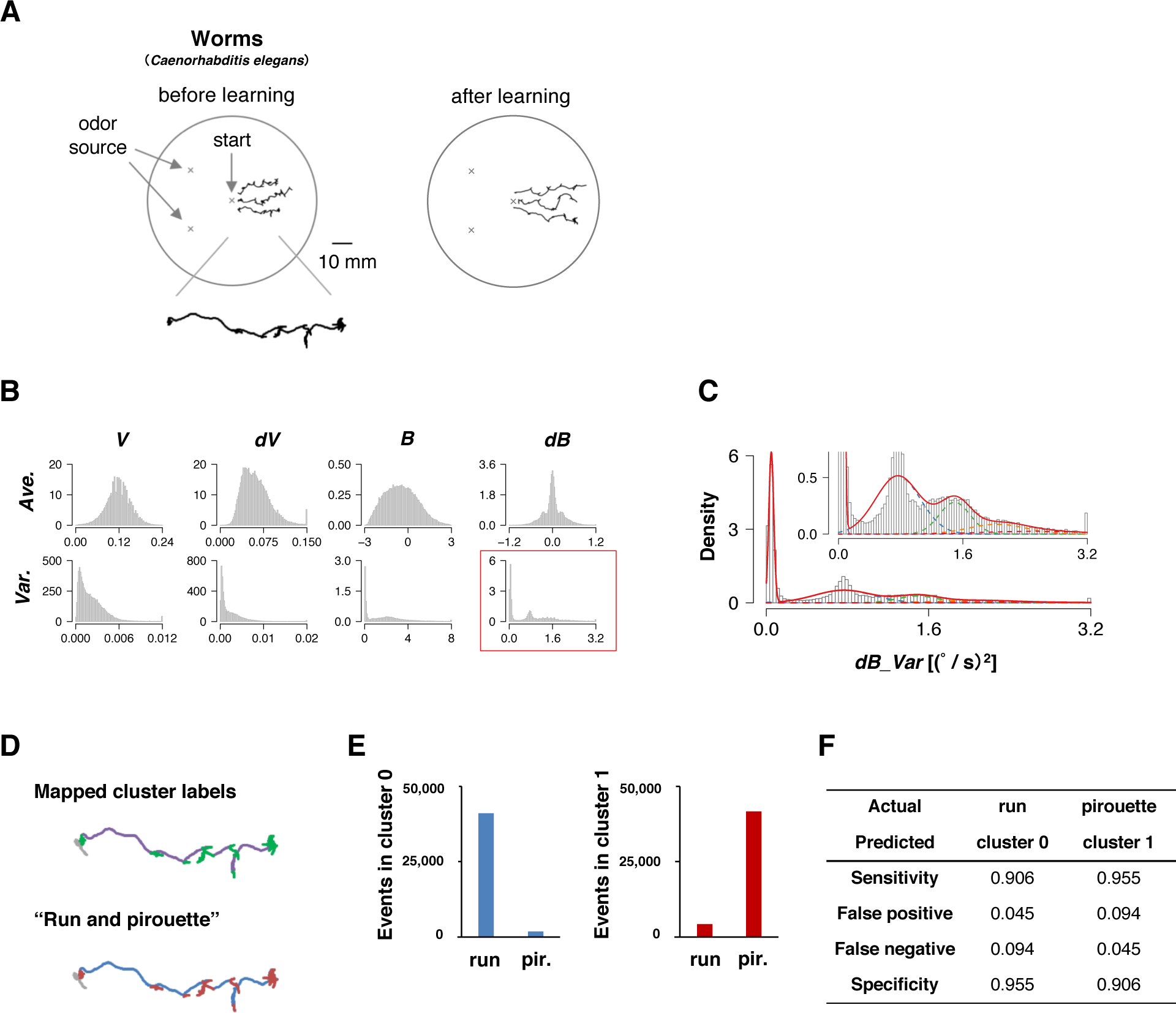
State estimation of worms. (A) Examples of the trajectories of 3 worms before (left) or after odor learning (right) in 12 min of 2-nonanone avoidance assay, overlaid on a schematic drawing of a 9 cm plate. One of the trajectories is magnified below. (B) The histograms of 8 basic behavioral features. Horizontal and vertical axes indicate the values and the density of each feature. *dB_Var* was chosen (red square) for the EM analysis. (C) Classification of *dB_Var* by the EM algorithm. Each cluster distribution (cluster 0, 1, 2, 3, and 4 are indicated by purple, blue, green, orange, and red dashed lines, respectively, similarly to the following figures) and the sum of clusters (red solid line) are shown. Inset is a magnified view. (D) Comparison of the cluster 0 and 1 (upper panel; purple and green, respectively) with run and pirouette (lower panel; blue and red, respectively) on a trajectory. The initial 2 minutes (gray in both panels) were excluded from the analyses because worms do not avoid the odor during the period (Kimura et al., 2010). (E) Event numbers of cluster 0 (left) and 1 (right) in run and pirouette. (F) Matching matrix of the state estimation.

We observed multiple peaks in the histogram corresponding to the variance values for bearing change (*dB_Var*, Fig. 2B). These values were automatically classified into 5 clusters by the EM algorithm (Fig. 2C). Upon mapping the clusters on to the trajectories, we found that cluster 0 corresponds to relatively straight part, while the other clusters correspond to more complex parts (Fig. 2D, upper panel).

Cluster 0 and clusters 1-4 mainly corresponded to “run” and “pirouette,” respectively, which are the two classical behavioral states of worms in various types of sensory behavior (Fig. 2D, lower panel) (Lockery, 2011; Pierce-Shimomura et al., 1999). “Run” constitutes relatively straight movement, while “pirouette” is characterized by short, straight movement divided by frequent large changes in angle (turns and reversals). Runs and pirouettes are usually classified based on a threshold value for the duration between consecutive large angle changes (Pierce-Shimomura et al., 1999), unlike in this method. We collectively regarded clusters 1-4 as “cluster 1” because the positional information of worms does not appropriately reflect their actual locations during pirouettes due to insufficient spatiotemporal resolution of the recording system for relatively long-distance navigation, such as odor avoidance behavior (Yamazoe-Umemoto et al., 2015). We found that more than 90% of the cluster 0 and 1 corresponded with the run and pirouette, respectively (Fig. 2E and F). Therefore, we concluded that the STEFTR method properly classified the odor avoidance behavior into distinct behavioral states.

Next, we applied the same process to the trajectories of penguins obtained using GPS devices. Penguins are good model wild animals for studying long-distance navigation given their relatively large body size for attaching GPS devices and their habit of returning to a colony, which makes the attachment and the recovery of GPS data easy (Yoda, 2018; Yoda et al., 2001). In this study, GPS and depth sensors were attached to 11 penguins from a colony in the Antarctic Continent; the depth sensors were used to evaluate the accuracy of state estimation (see below). The penguins moved by walking and swimming about 10 km for feeding, and each dataset contained up to 2 days of data recordings (Table 1 and Fig. 3A). Like in the case of worms, 8 basic behavioral features were extracted from the penguin trajectory data and represented as histograms. We chose the average velocity (*V_Ave*) as it showed multiple peaks in the histogram. The EM algorithm classified it into 5 clusters (Fig. 3B lower panel, C and D upper panel).

**Figure 3.**
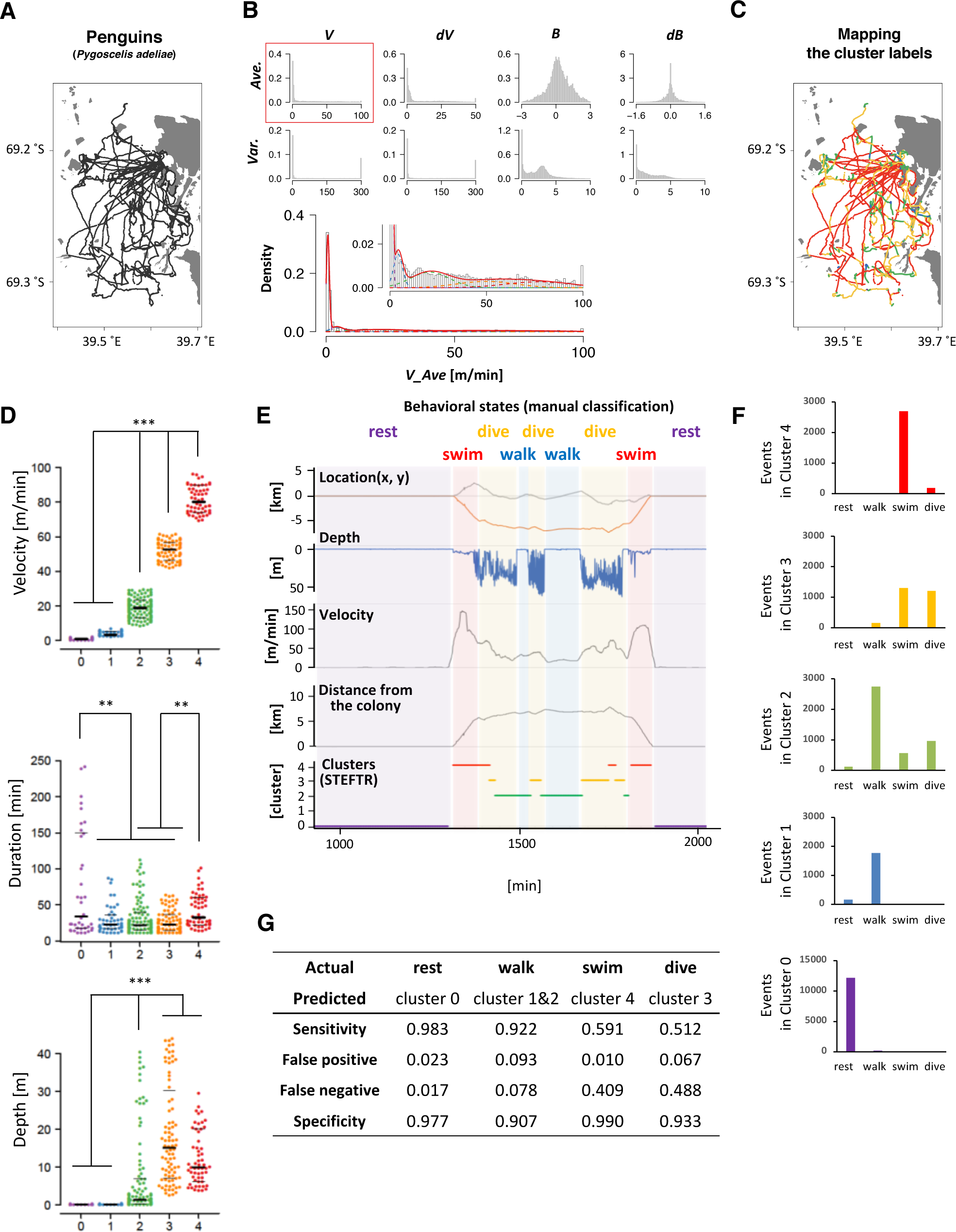
State estimation of penguins. (A) The trajectories of 11 penguins (black lines) on the Antarctic Continent (gray area; white area is the sea). Horizontal and vertical axes indicate longitude and latitude, respectively. (B) The histograms of 8 basic behavioral features (upper panels) and the classification by the EM algorithm. *V_Ave* was chosen (red square) and classified into 5 clusters (lower panels). (C) Mapping of the five clusters on the trajectory. (D) Differences in velocity, duration, and depth among the clusters. Each dot represents a cluster bout, and the bars represent the median and the first and third quartiles. Significant differences among clusters suggest that the clusters correspond to different behavioral states. Statistical values were calculated using Kruskal-Wallis test with *post hoc* Dunn’s test. \*\**p* < 0.01, \*\*\**p* < 0.001. (E) An example of comparison of the clusters from the STEFTR analysis with the behavioral states by manually classified labels, which is based on diving depth, movement speed recorded from GPS data, and distance from the colony. (F) Event numbers of each clusters. (G) Matching matrix of the state estimation. The statistical details are described in Supplementary Table 1.

Interestingly, the clusters exhibited significantly different distributions in multiple behavioral features. For example, the values for the duration of each bout were much longer in cluster 0 than in clusters 1, 2, and 3 (Fig. 3D, middle panel). In addition, although the clusters were classified only based on the horizontal velocities, the depths for cluster 0 and 1 were significantly closer to zero than those for clusters 2, 3, and 4 (Fig. 3D, lower panel). These results are consistent with the idea that each cluster reflects a behavioral state that is a complex function of multiple behavioral features.

To evaluate whether the clusters actually reflect different behavioral states, we manually classified the penguin behavior into four states (resting, transit by walking, transit by swimming, and diving) based on diving depth (from depth sensor), movement velocity and distance from the colony (both calculated from the GPS positional information) (Yoda et al., 2001). Penguins stayed and rested at the colony (location does not change much; depth is zero), moved on land and ice mainly by walking (location changes relatively slowly; depth is near-zero), swim in the sea to go to the foraging area (location changes quickly; depth is relatively shallow in general and sometimes increases when they are moving towards the feeding area), dive deeply at the foraging area (location does not change much; dives occur continuously in bouts), and then come back to the colony by swimming and walking. The resting at the colony and swimming correlated with clusters 0 and 4, respectively (Fig. 3E and F). In addition, most of clusters 1 and 2 correlated with walking, while about 50% of cluster 3 corresponded to diving (Fig. 3F). Thus, when a behavioral state is classified to a cluster other than cluster 3, the penguin is likely to be resting, walking, or swimming. If a behavioral state is classified to cluster 3, which is ~10% of all the behavior recorded, the penguin is either diving or swimming. Remarkably, although the clustering is only based on the trajectories of 11 penguins for a few days, the false positive rates were less than 10% and the sensitivity of the analysis was greater than 90% in all the cases (Fig. 3G). Thus, we concluded that the STEFTR method can reasonably estimate different behavioral states only based on trajectory data.

The STEFTR method was also applied to the trajectories of flying seabirds in the wild and rats in the laboratory. The seabirds, *Calonectris leucomelas,* traveled ~100 times longer distances (up to 1,000 km) with ~10 times the speeds compared to penguins (Matsumoto et al., 2017; Yoda et al., 2014). Like in the case of penguins, the average velocity (*V_Ave*) showed multiple peaks and was classified into 5 clusters (Fig. 4A and B) with significantly different profiles for duration and directedness (Fig. 4C). In the case of rats, the variance of velocity change (*dV_Var*) was chosen, and the EM algorithm classified the behavior into 4 clusters (Fig. 4D and E). Interestingly, significant differences were observed in several features such as duration and speed (Fig. 4F). Such information can help ecologists estimate the candidates for feeding areas where fishes may be more densely distributed and discover biologically important marine areas. It can also help neuroscientists in estimating candidate conditions to further explore specific neural activities.

**Figure 4.**
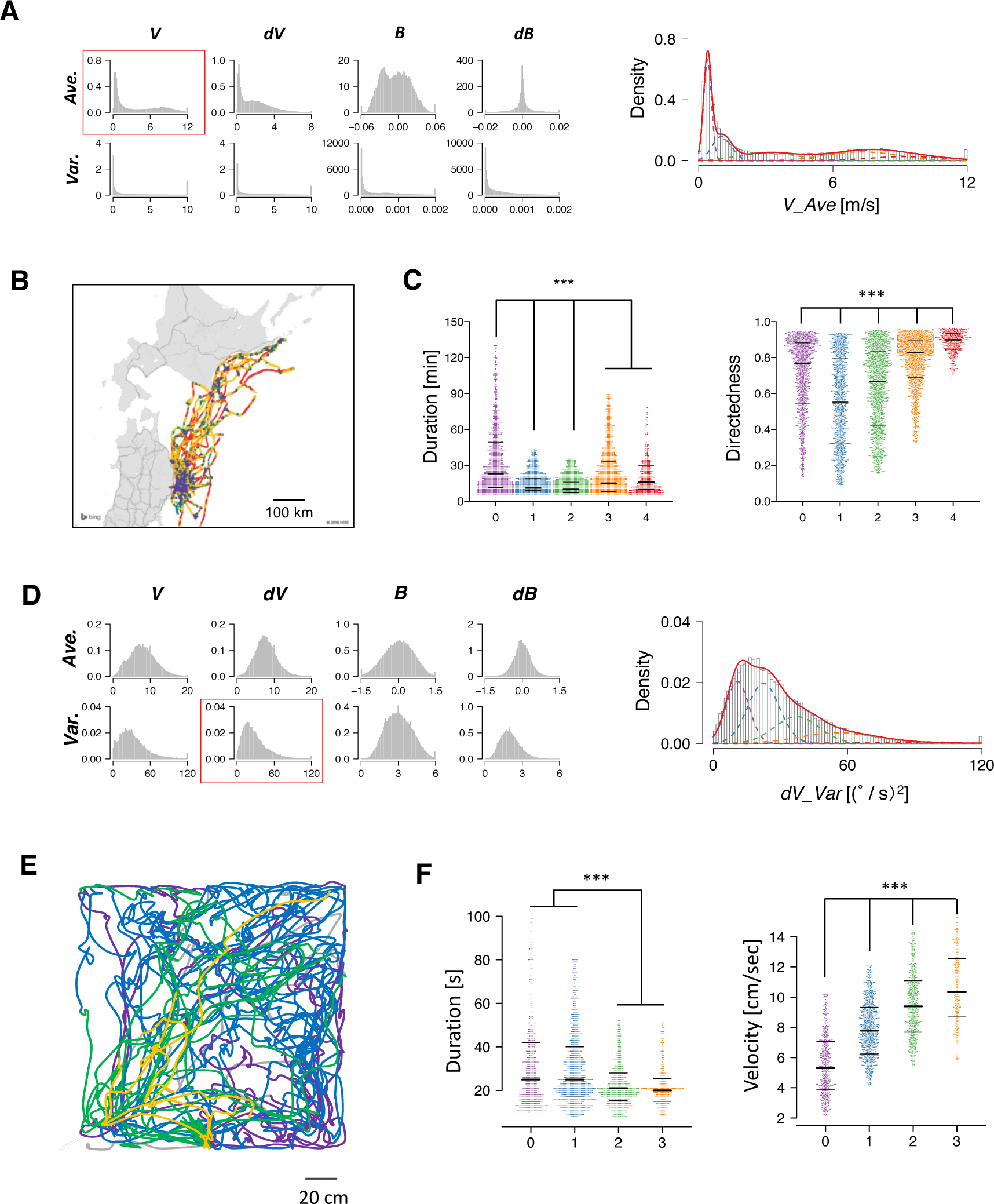
State estimation of flying seabirds in the Pacific Ocean (A-C) and rats in the open maze (D-F). (A) From the histograms of 8 basic features of flying seabirds (left panels), the average of velocity (*V_Ave*) was chosen and classified into 5 clusters (right panel). (B) Mapping of the clusters on the trajectory. The gray region is the northern part of Japan (Tohoku and Hokkaido area), while the white region is the sea. (C) Significant differences were observed in duration (left) and directedness (right). (D) From the histograms of 8 basic features of rats (left panels), the variance of velocity change (d*V_Var*) was chosen and classified into 4 clusters (right panel). (E) An example of trajectories of one rat. (F) Significant differences were observed in duration (left) and velocity (right). Each dot represents a cluster bout, and the bars represent the median and the first and third quartiles. Statistical values were calculated using Kruskal-Wallis test with *post hoc* Dunn’s test. \*\*\**p* < 0.001. The statistical details are described in Supplementary Table 1.

### Comprehensive extraction of behavioral features modulated by learning

As a second part of the STEFTR method, comprehensive feature extraction was performed by comparing a specific behavioral state in two different conditions, such as cluster 0 of worms before and after learning. Comprehensive semi-automated analysis can be very helpful to compare behavioral features in two conditions. This is because even when the overall result of behavioral responses are different in two conditions, it is still difficult to quantitatively determine which part of the trajectories are different (Fig. 2A left and right, for example). Furthermore, even if several behavioral features are found to be different, it is possible that other more prominent feature differences may exist. We considered that learning-dependent changes in behavior should be one of the best models for comprehensive feature extraction because the differences in behavioral features should reflect learning-dependent changes in neural/brain activities.

As an useful index for feature extraction, we chose information gain, the index for decision tree analysis (Quinlan, 1986). Binary decision tree analysis is for splitting a dataset into two sub-groups by automatically selecting the best feature and its parameter showing the largest information gain (i.e., difference of uncertainty, or “information entropy”, between before and after division). Each data point is then classified into one of the sub-classes based on whether it has a larger or smaller value than an automatically determined threshold. When applied for binary classification, decision tree analysis automatically evaluates the classification performance of a large number of features as designed by the researchers. This analysis results in the extraction of certain features, allowing us to easily understand the utility of particular features in the classification. This approach is substantially different from those that employ support vector machines and/or deep neural networks, wherein the relationships between the classification and the features of the data cannot be easily discerned.

We first analyzed learning-dependent changes in worm odor avoidance behavior. Worm odor avoidance behavior is enhanced by pre-exposure to the odor as a type of non-associative learning, and pre-exposed worms migrate significantly longer distances from the odor source than control worms do during the same period (Fig. 2A) (Kimura et al., 2010). This phenomenon is interesting because prior exposure to a stimulus generally causes reduction, instead of enhancement, of the response to the stimulus through adaptation or habituation. Although this is a simple form of learning, this odor learning is modulated by multiple neuromodulators, including dopamine, octopamine (the worm counterpart of mammalian noradrenaline), and neuropeptides (Kimura et al., 2010; Yamazoe-Umemoto et al., 2015). Previous quantitative analyses have shown that the enhanced odor avoidance behavior is not caused by changes in speed, but mostly by increases in run duration (Yamazoe-Umemoto et al., 2015). However, this did not rule out the possibility that other behavioral features play more profound effects.

As an example of comprehensive feature extraction from a behavioral state, we focused on learning-dependent changes in cluster 0 (run) because the values of their centroid migration are quantitatively more reliable than cluster 1 (pirouette) as mentioned above. In addition to the basic behavioral features used for the estimation of behavioral states (*V*, *dV*, *B*, and *dB*), we also calculated directedness (*Dir*) (Gorelik and Gautreau, 2014), and the odor concentration (*C*) and temporal change in odor concentration (*dC*) that each worm experienced during the odor avoidance behavior; *C* and *dC* were calculated based on actual measurements of the dynamic odor gradient (Tanimoto et al., 2017; Yamazoe-Umemoto et al., 2018). For these, we calculated the initiation (*Ini*), middle (*Mid*), termination (*Ter*), and all (*All*) values of a cluster 0 segment (Fig. 5A). In addition, different time windows (1-6 s in this case) were used to calculate these values because a behavioral feature could be apparent only within a specific temporal window (for example, velocity of run (i.e. cluster 0) starts decreasing 2 s before the end of a run (Pierce-Shimomura et al., 1999)). We also calculated durations (*Dur*) of cluster 0 and 1, and the weathervane index (*WV*) (Iino and Yoshida, 2009). Information gain for each of these features were compared between naive/mock and pre-exposed conditions (Fig. 5B, for example). The information gain values for each of the features have been summarized in Table 2, and the details are described in Supplementary Tables 2-11.

**Table 2.**
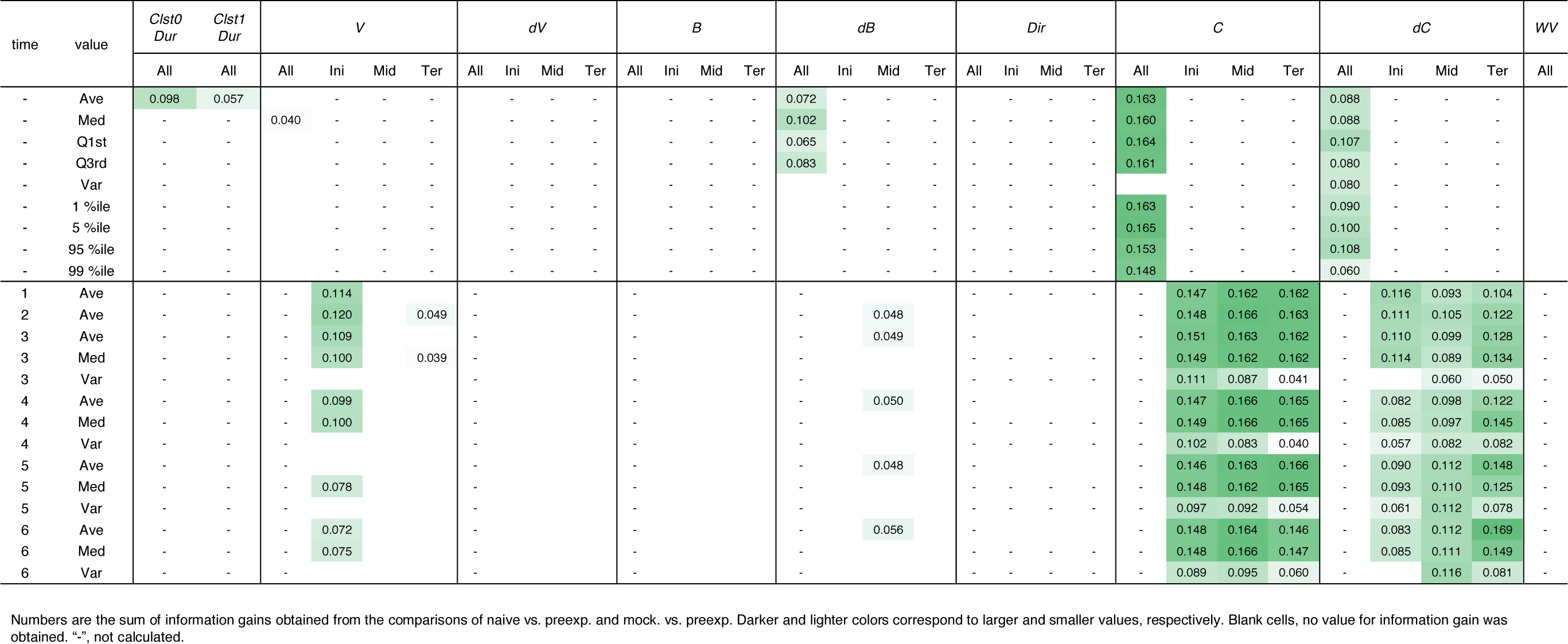
Learning-dependent features extracted from cluster 0 in odor avoidance behavior of wild-type worms.

**Figure 5.**
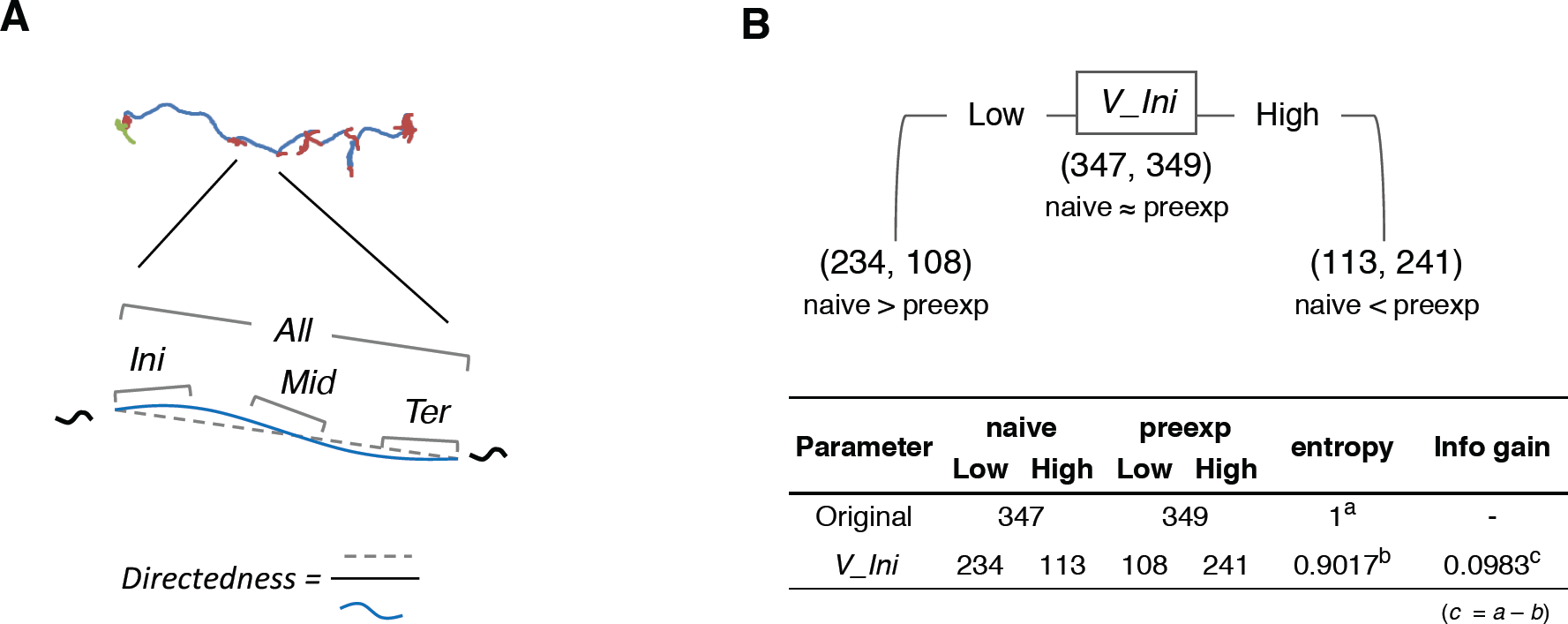
Feature extraction of worm behavior. (A) Schematic drawing of the behavioral features. (B) One example (*V_Ini*; average of time window 2) of calculation of information gain.

Through this analysis, we were able to find new as well as previously known behavioral features that exhibited learning-dependent changes. First, we found that the duration of each cluster 0 (*Clst0Dur*) exhibited higher information gain (Table 2), which corresponded to significantly increased cluster 0 duration (Fig. 6B). This result is consistent with the findings from previous reports (Kimura et al., 2010; Yamazoe-Umemoto et al., 2015), highlighting the reliability of this method. We also found that the velocity at the beginning of each cluster 0 (*V_Ini*) consistently exhibited higher information gain in the average and median values in multiple time windows (Table 2); these values were also significantly different in the pre-exposed worms as compared to the control worms (Fig. 6C). The previous study has not identified this difference as only average values per run (*i.e.* cluster 0) have been calculated in the study (Yamazoe-Umemoto et al., 2015). Although the contribution of this behavioral feature to enhanced odor avoidance is unclear at present, our results indicate that the STEFTR method can reveal characteristic feature(s) under specific conditions, which is difficult for human analyses to accomplish.

**Figure 6.**
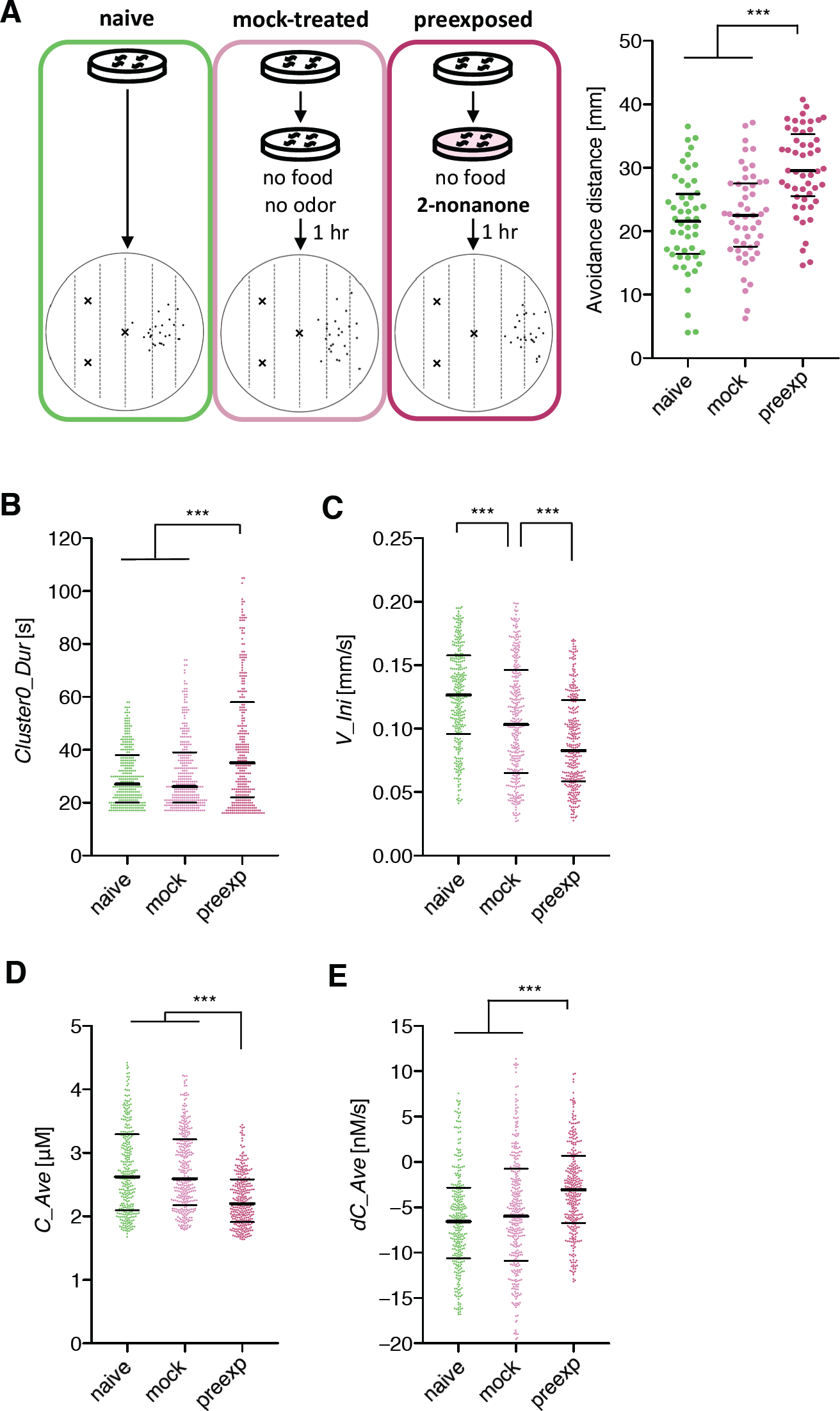
Extracted features modulated by the odor learning. (A) Enhanced odor avoidance behavior in worms caused by odor pre-exposure. Left: End-points of 25 worms in each condition plotted on a schematic representation of the assay plate. Right: Avoidance distance (distance between the center line of the plate and the end-point of the behavior) of each worm. Each dot represents a worm. Significant differences were observed between the pre-exposed worms and the naive and mock-treated worms (\*\*\**p* < 0.001, Kruskal-Wallis test with *post-hoc* Dunn’s test). (B, C, D and E) Distributions of extracted features. Duration (B), the initial value of velocity (C; average of time window 2), the average odor concentration (D), and the average odor concentration change (E) of each run (\*\*\**p* < 0.001, Kruskal-Wallis test with *post-hoc* Dunn’s test). Each dot represents a cluster bout, and the bars represent the median and the first and third quartiles. The statistical details are described in Supplementary Table 1.

Odor stimuli during runs, which likely drive the worms’ odor avoidance behavior, were also found to be consistently modulated (Table 2). In fact, odor concentration (*C*) was significantly lower, and the temporal change in odor concentration (*dC*) was significantly closer to zero (i.e. shallower) in a learning-dependent manner (Fig. 6D and E). Because the previous study demonstrated that worm odor avoidance behavior depends on *dC* rather than *C* at least in the naive condition (Tanimoto et al., 2017), one possibility is that the changes in the responsiveness of worms to *dC* is the underlying reason for the enhanced odor avoidance. However, it is also possible that the odor-experienced worms were somehow located farther away from the odor source than the unexperienced worms, and hence, sensed lower odor concentrations and shallower odor concentration change than the latter.

### Responsiveness of sensory neurons to odor increase was modulated by the odor learning

If the change in sensitivity to *dC/dt* is the reason underlying enhanced odor avoidance behavior, it should be associated with changes in neural activity. Thus, we analyzed the responsiveness of a likely candidate, ASH nociceptive neurons (Bargmann, 2006; Kaplan, 1996). Previously, we have established the OSB2 microscope system that allows for *in vivo* calcium imaging of *C. elegans* neurons in the presence of odor stimuli resembling those that the worms experience during the odor avoidance behavior in the plates (Fig. 7A) (Tanimoto et al., 2017). Using the OSB2 system, we found that ASH neurons are the major sensory neurons to cause pirouettes upon increases in 2-nonanone concentration (Tanimoto et al., 2017). However, whether the ASH response is modulated by 2-nonanone experience has not yet been studied.

We found that ASH responses were indeed modulated by prior odor experience. When the worms were stimulated with a 5 nM/s odor increase rate, which is the lowest rate of change to cause the threshold-level behavioral response in the previous study (Tanimoto et al., 2017), ASH neurons in naive as well as mock-treated worms exhibited robust responses (Fig. 7B). However, the ASH responses were significantly reduced in the pre-exposed worms (Fig. 7B and C). This suggests that prior odor experience causes a reduction in the neuronal response to a slight increase in odor concentration, subsequently causing longer run durations and enhanced odor avoidance behavior (Fig. 7D).

**Figure 7.**
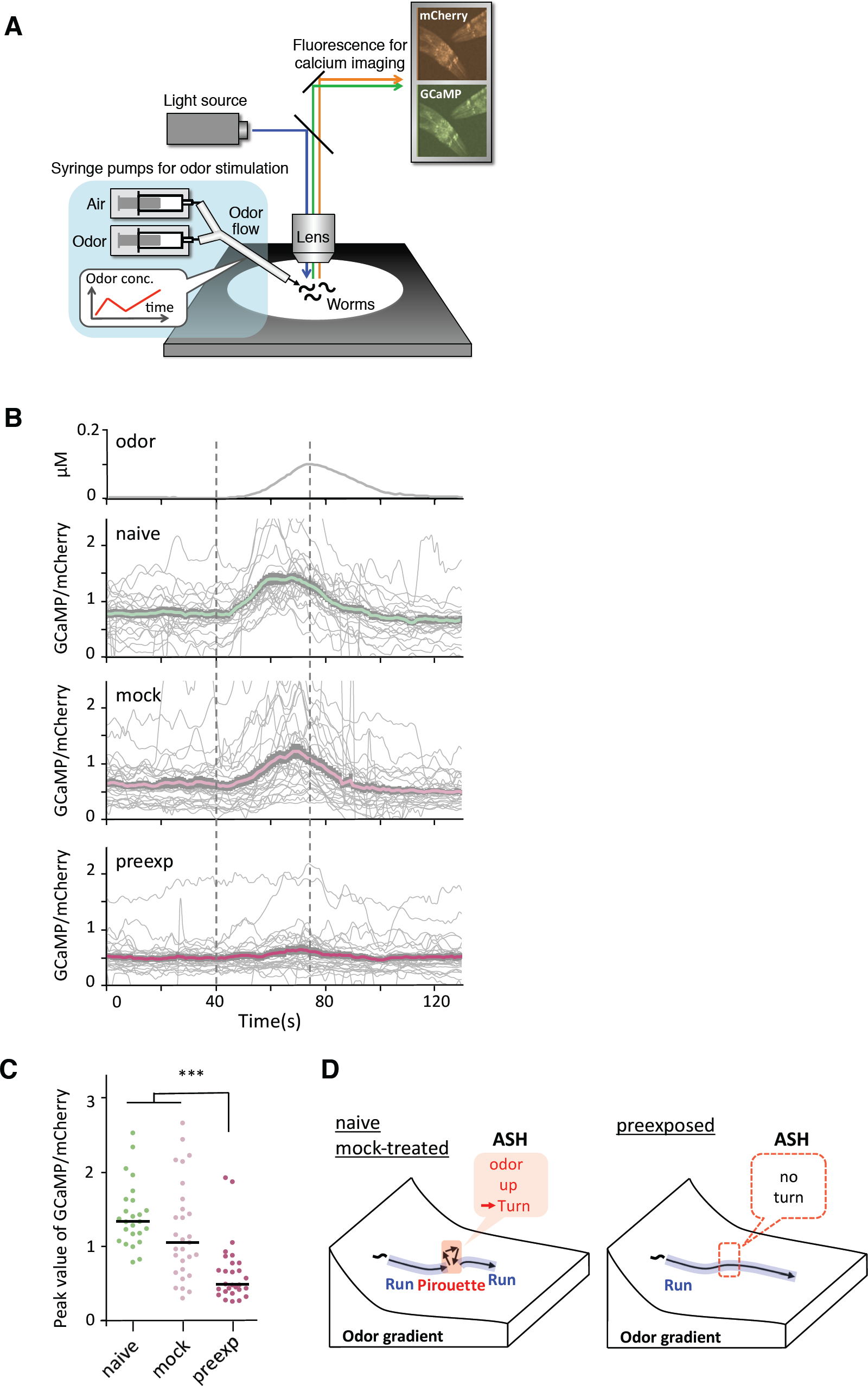
Sensory responses to slight increases in odor concentration were reduced by pre-exposure to the odor. (A) A schematic drawing of calcium imaging of neural activity of worms under odor stimuli. Several immobilized worms were simultaneously exposed to an odor flow whose concentration was changed by controlling syringe pumps. (B) Responses (GCaMP/mCherry) of ASH neurons in naive (n = 25), mock-treated (n = 29), and pre-exposed (n = 26) worms. Thick lines with gray shadows indicate mean ± standard error of mean, while thin lines indicate individual responses. (C) Distributions of peak values during the odor-increasing phase (*t* = 40-80 s) shown in panel B. The bars represent the median. (\*\*\**p* < 0.001, Kruskal-Wallis test with *post-hoc* Dunn’s test). (D) A model relationship between odor concentration change and behavioral response during navigation along the odor gradient. When naive and mock-treated worms sensed a slight increase in the odor concentration, which is a sign of migrating in the wrong direction, they stopped a run and started a pirouette to search for a new direction. In contrast, the pre-exposed worms did not respond to a slight increase in odor concentration, leading to longer run durations (and shorter pirouette durations in total as a consequence), which likely contribute to the enhanced avoidance distance. The statistical details are described in Supplementary Table 1.

### Extracted behavioral features of mutant strains correspond to gene function

Next, we comprehensively analyzed learning-dependent behavioral changes in the mutant *C. elegans* strains. Many mutant strains of *C. elegans* showing impaired learning have already been isolated and characterized (Bargmann, 2006; Sasakura and Mori, 2013), and the behavioral abnormalities observed in these mutants should reflect the role of the causal genes in neural function. In fact, we have previously shown that two different genes involved in the enhanced odor avoidance behavior cause different abnormalities in behavioral features when mutated (Yamazoe-Umemoto et al., 2015). However, as the behavioral features exhibited by a mutant strain could be different from one another, identification of abnormal behavioral features is often laborious and time-consuming.

In addition to studying the previously described mutants with defective enhanced odor avoidance behavior (*egl-3* and *egl-21* for neuropeptide biosynthesis, and *dop-3* for dopamine receptor) (Kass et al., 2001; Suo et al., 2004; Yamazoe-Umemoto et al., 2015), we also analyzed mutant strains found to be involved in the phenomenon in this study (*ocr-2* and *osm-9* for TRP channels; *tax-4* for CNG channel; *eat-4* for vesicular glutamate transporter; *pkc-1* for protein kinase) (Colbert et al., 1997; Komatsu et al., 1996; Land et al., 1994; Lee et al., 1999; Tobin et al., 2002) (Table 3).

**Table 3.**
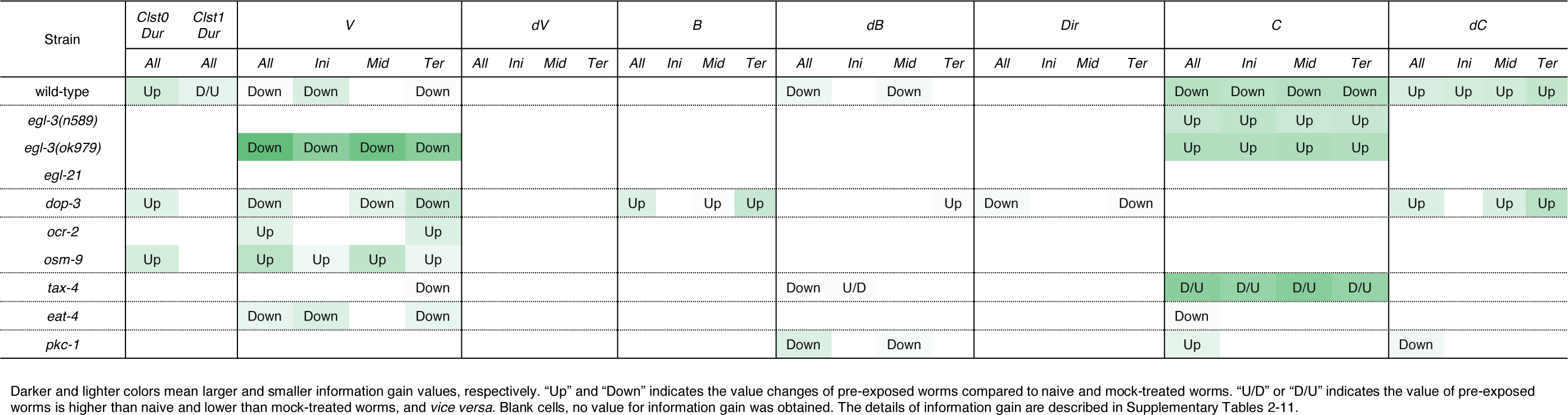
Patterns of learning-dependent behavioral features of cluster 0 in odor avoidance behavior of mutant worms.

Neuropeptide mutant strains did not exhibit learning-dependent changes in behavioral features, except for the velocity of *egl-3(ok979)*. This result is consistent with the previous finding that neuropeptide signaling is required for the acquisition of odor memory (Yamazoe-Umemoto et al., 2015). *egl-3(ok979)* may have exhibited stronger phenotypes than *egl-3(n589)* because they are nonsense and missense mutants, respectively. Also consistent with the previous report (Yamazoe-Umemoto et al., 2015), the *dop-3* mutants exhibited abnormalities in direction-related behavioral features (*B* and *Dir*) while the changes in cluster 0 durations and velocities are similar to those of wild-type worms (Table 3). Furthermore, with respect to the newly added mutant strain, similar patterns are observed in *ocr-2* and *osm-9* mutants of the TRP channel involved in sensory perception. On the other hand, *tax-4*, which is also involved in sensory perception but expressed in a different set of sensory neurons (Komatsu et al., 1996; Tobin et al., 2002), and *eat-4* and *pkc-1* mutants showed different patterns of abnormalities. Taken together, our results suggest that the patterns of features extracted from mutant strains may reflect functional groupings of the mutated genes. Thus, profiling and classification of extracted mutant features of unknown genes may be useful in the estimation of their physiological functions.

### Feature extraction of fly sexual behavior

Next, we applied the technique to comprehensive feature extraction of animal behavior under specific situations in two different conditions—heterosexual chasing behavior of *Drosophila melanogaster* with or without pheromone sensation. On an experimental tracking system (Fig. 8A), male flies chased the target female flies’ abdomens after tapping them with their forelegs to sense the cuticular pheromone, although males do not show such chasing behavior before tapping (Kohatsu et al., 2011). While this pheromone-driven behavior has been generally used for the observation of neural activity during courtship behavior in fruit flies, the behavioral features have not yet been elucidated in great detail.

**Figure 8.**
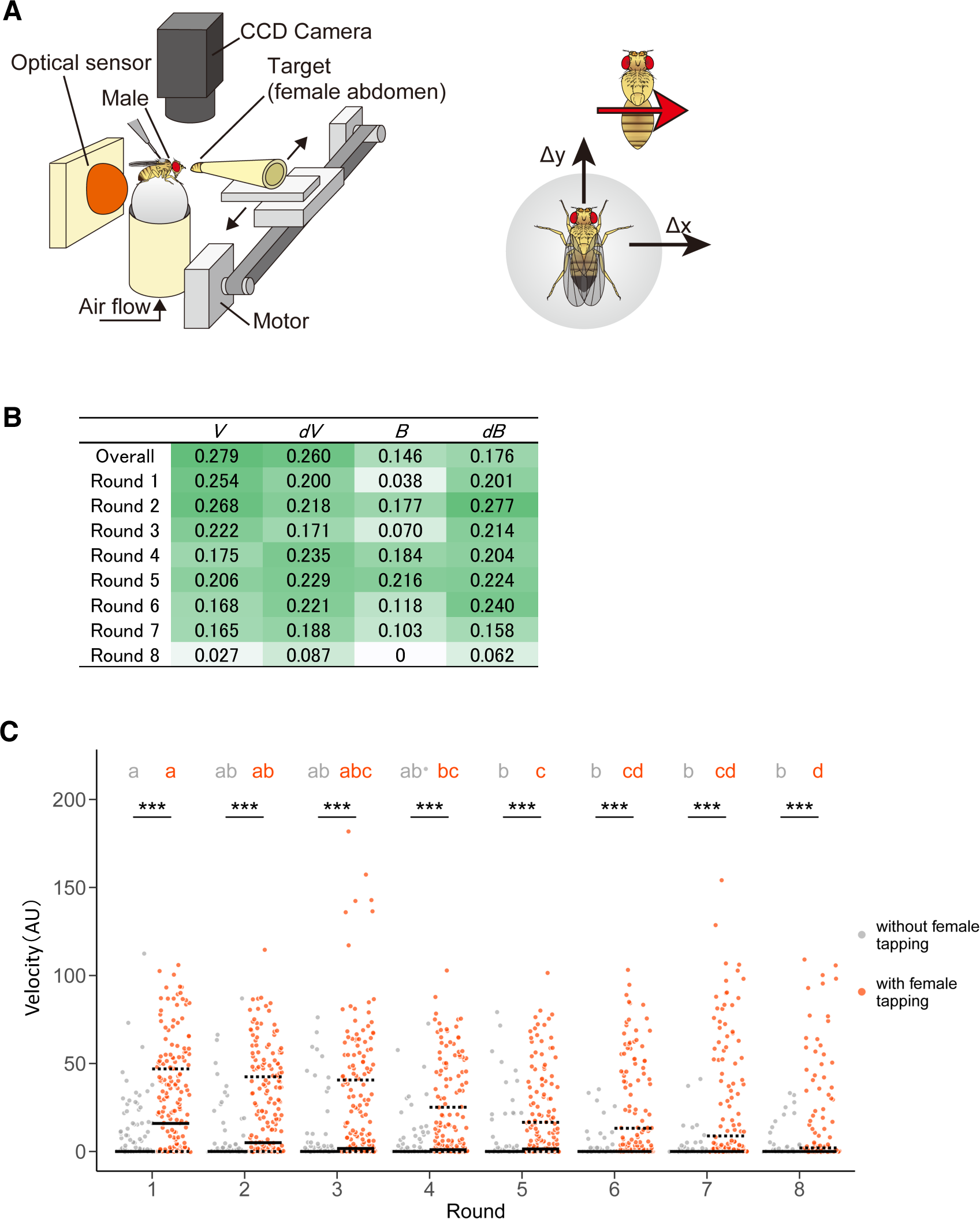
Pheromone-driven responses of male fruit flies decreased over time. (A) A schematic drawing of the experimental setup. A female fly was actuated leftward and rightward in front of the male fly. The locomotion of the male fly was monitored by an optical sensor, which recorded lateral (Δx) and forward (Δy) movements at 4 Hz. (B) Information gain. Darker and lighter colors mean larger and smaller values, respectively. (C) Distribution of velocity in the chasing behavior of male flies. Control (without female tapping, gray dots) and experimental (with female tapping, orange dots) groups are shown. Solid and dotted lines represent the median and the first and third quartiles, respectively. Asterisks indicate the statistical significance between the control and test groups (Mann-Whitney U test followed by Bonferroni test for multiple comparison correction, p < 0.05). Different characters in each group indicate statistical significance among rounds (Steel-Dwass test, *p* < 0.05). The statistical details are described in Supplementary Tables 12 and 13.

In this study, we used a tracking system as described in previous studies, where a male was exhibited the moving female abdomen with 8 time left-right round trip after a pheromone sensation (Fig. 8A) (Kohatsu et al., 2011). We detected positive information gains in the velocity, changes in velocity, bearing, and changes in bearing in a pheromone sensation-dependent manner (Fig. 8B). Unexpectedly, the information gains of velocity were higher in the earlier round trips and decreased over the trips (Fig. 8B). This suggests that the pheromonal effect promoting chasing behavior decreases over time. To confirm the result of the STEFTR method, we re-analyzed the speed of the male locomotion along with time-series. Consistent with the STEFTR result, the velocity of males that had tapped the female significantly decreased over the trips (orange group in Fig. 8C; significant differences between rounds 1, 4, and 8), whereas that of control flies (gray group in Fig. 8C) remained mostly unchanged (significant difference only between rounds 1 and 5). Thus, the STEFTR method can even uncover behavioral features that fluctuate over time. The decreased tracking velocity may reflect a decrease in motivation in the fly brain, which can be assessed directly by observing the temporal changes in neuronal activity related to the courtship-motivation circuit in the fly brain (Yamamoto and Koganezawa, 2013; Zhang et al., 2018).

### Feature extraction of learning-dependent modulation of acoustic navigation in bats

To further demonstrate the general applicability of the method, we examined features of acoustic navigation in bats. We have previously reported that bats improve their flight trajectory in an indoor space with obstacles in a learning-dependent manner (Yamada et al., in revision). Here, we optimized features such as velocity (*V*), distance to the obstacle chain array (*R_obs* and *R_x*), and horizontal bearing of the flight (*B_hori*) for the experimental paradigm (Fig. 9A and B). Interestingly, although the velocity (*V*) itself was modulated by flight experience, the change in velocity (*dV*) was not (Fig. 9C and D), suggesting that bats determine flight speed before initiating navigation, but not during navigation, at least in this experimental conditions. As the vocalizations of bats reflect their attention or decisions (Moss and Surlykke, 2010), our results suggest that the STEFTR method can be used to elucidate such higher brain functions during navigation.

**Figure 9.**
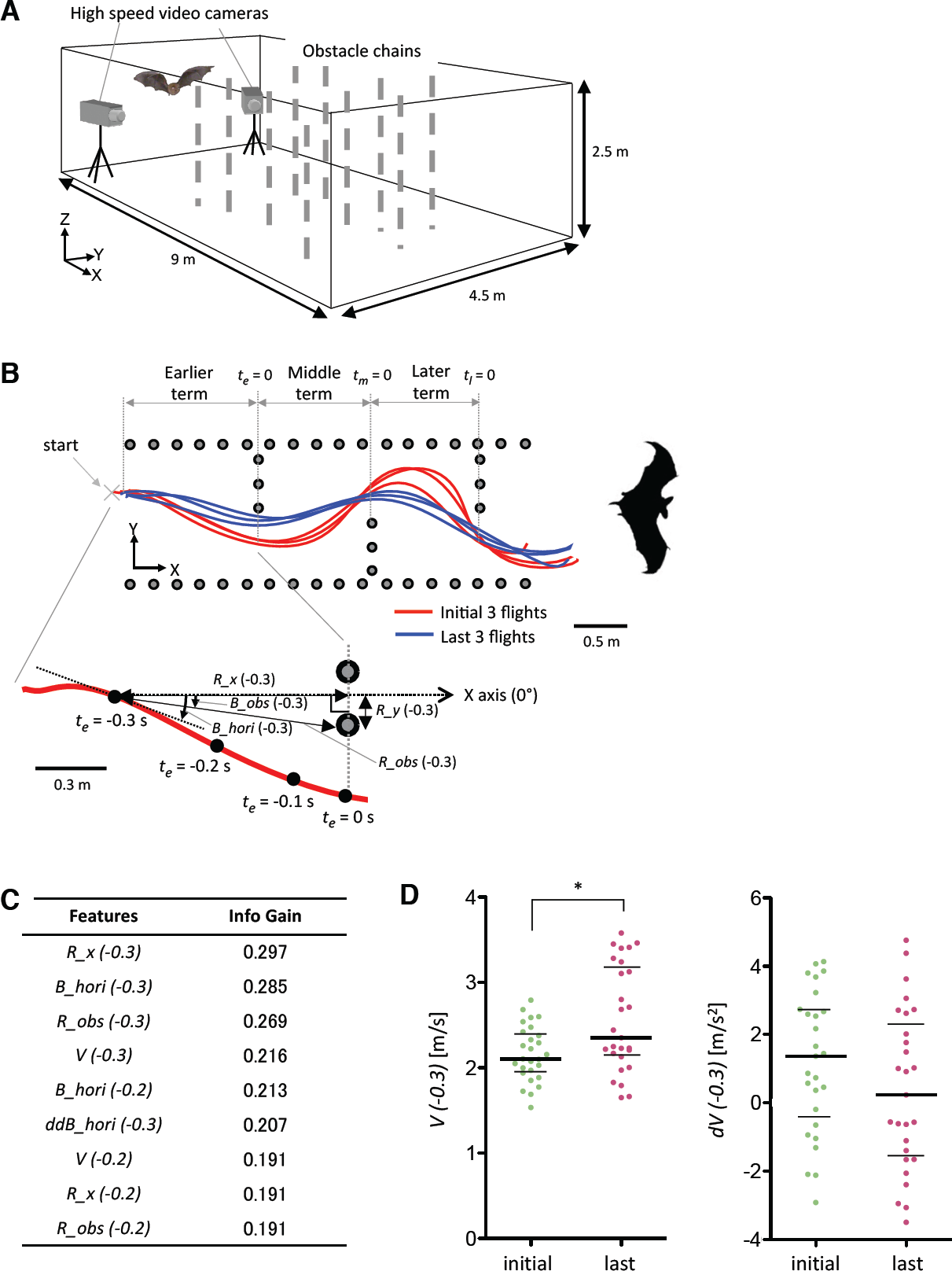
Learning-dependent changes in bat acoustic navigation. (A) The experimental setup for monitoring the 3D flight trajectory of a bat during obstacle avoidance flight in a chamber. (B) Representative flight trajectories of a bat in the horizontal plane during repeated flights in the obstacle course. The figure on the top combines the first three (red) and last three (blue) flight trajectories. Each behavioral feature was collected in three segments: earlier, middle, and later terms. The figure on the bottom shows an expanded view of the earlier term in the first flight. Definition of the horizontal bearing of the flight (*B_hori*), distance (*R_obs*), and bearing (*B_obs*) of the bat to the nearest edge point of the obstacle chain array, longitudinal directional distance to the frontal chain array (*R_x*), and lateral directional distance to the inside pitch of the chains array (*R_y*) are indicated. Time windows for the analysis of each behavioral feature were 0.1, 0.2, or 0.3 s before or while (*t* = 0) passing through the chain array. (C) A list of extracted features of bat acoustic navigation modulated by flight experience. (D) Distributions of *V(−0.3)* and *dV(−0.3)* are plotted. The bars represent the median and the first and third quartiles. (\**p* < 0.05, Kruskal-Wallis test with *post-hoc* Dunn’s test). The statistical details are described in Supplementary Table 1.

## DISCUSSION

Measuring and analyzing behavior is one of the most prominent steps in understanding brain function. In order to utilize “behavioral big data”, we developed a hybrid supervised/unsupervised technique, the STEFTR method, to estimate behavioral states and to efficiently extract behavioral features solely from the trajectories of animal movement. The behavioral states of worms and penguins estimated with the STEFTR method were in reasonable agreement with the ones based on previous knowledge, highlighting the validity of our method. In addition, one of the learning-dependent behavioral features extracted from worms corresponded to a change in neural activity. Furthermore, we were able identify temporally dynamic changes through feature extraction from fly courtship behavioral data.

One of the advantages of the STEFTR method is its versatility. Multiple methods have been reported for the behavioral analysis of specific animals under specific conditions (Baek et al., 2002; Branson et al., 2009; Brown et al., 2013; Dankert et al., 2009; Kabra et al., 2013; Mathis et al., 2018; Robie et al., 2017; Stephens et al., 2008; Vogelstein et al., 2014; Wiltschko et al., 2015). However, the animals and experimental conditions for which each of these methods can be applied are rather limited. For example, it is still not easy to effectively and robustly extract an animal’s posture from a video image, especially in the wild. Even in laboratories, the parameters generally need to be adjusted again when imaging conditions changed (Dell et al., 2014; Egnor and Branson, 2016). In contrast to these methods, our method allows for behavioral analysis based on positional information that can be extracted from animal video images as well as from different methods such as GPS device.

As a first step of the STEFTR method, we estimated behavioral states from animal trajectories. State estimation is one of the critical processes of movement analysis of animals in the wild as well as of cars and people in ecology and data science, respectively (Egnor and Branson, 2016; Jonsen et al., 2013; Patterson et al., 2008; Zheng, 2015). However, the analytical methods that can be applied to the analysis of various types of animals (and cars and people) are still in debate (Gurarie et al., 2016; Zheng, 2015). In the STEFTR method, we aimed to analyze behavior without previous knowledge of the animal and/or experimental condition and independent of the spatiotemporal scale of the behavior. We analyzed migratory velocity and direction, the most fundamental elements of moving objects, with appropriate moving-averaged data. Interestingly, in case of both worms and penguins, the moving average with a temporal window that is about 1% of median length of the behavioral record provided results that were reasonably consistent with behavioral states categorized by manual labeling (Fig. 2D-F and 3E-G). This result suggests that the ~1% temporal window works as a low-pass filter for properly estimating behavioral states in a scale-free manner. We estimated the number of behavioral states to be equal to or less than 5 (Jonsen et al., 2013), which worked well in the above-mentioned cases.

State estimation based only on trajectory analysis using the STEFTR method is not perfect, as shown in the case of penguins (Fig. 3E). However, the estimated behavioral states likely provide us with important information for further experiments, such as when and where in the spatiotemporal behavioral profile of the animal behavior should be analyzed in detail, especially in the case where the behavior has not been studied intensively in quantitative manner. For example, relatively small movements at places distant from their nest in the wild may correspond to the feeding area. For neurobiological/physiological analysis, the transition from one state to the other could be triggered by a specific change in the sensory stimulus and associated with specific neural activities. It should also be noted that the STEFTR method allows semi-automatic state estimation and feature extraction, which is suitable for large-scale behavioral analysis of mutant strains of laboratory model animals.

The Hidden Markov Model (HMM) has been used for the estimation of behavioral states (Gurarie et al., 2016; Jonsen et al., 2013). However, the HMM depends on the idea of the Markov process, wherein the state transition is probabilistic. In general, state transition in animal behavior is triggered by external as well as internal stimuli, which are not always probabilistic. While the STEFTR method assumes a probabilistic normal distribution of a behavioral feature in a state, it does not assume probabilistic transition between states.

For comprehensive feature extraction, we used information gain, an index used in decision tree analysis. Decision tree analysis is one of the machine learning techniques used for classification. Classification analysis involves classifying new, unlabeled data into appropriate classes using characteristic features and the parameters that have been extracted from the known class-labeled data. In the present study, however, the classification itself was not meaningful because the data were already classified (with/without learning or with/without sex pheromone). Instead, we focused on the procedure in the classification that identifies features useful for distinguishing between the two classes. In other words, behavioral features that are different between two classes (*i.e.,* conditions) should be able to effectively classify the behavioral data of animals in two conditions.

Although the STEFTR method does not directly provide information about brain/neural activity underlying animal behavior, it provides us with clues required to formulate hypotheses related to the experimental investigation of the neural activity, as shown in the case of learning-dependent changes in behavioral features and neural activity (Fig. 7). For example, animals in the wild experience continuously changing visual, auditory, and olfactory stimuli, each of which contain multi-dimensional information (color, shape, tone, different chemical compounds, etc.). Therefore, it is difficult to identify which aspect(s) of the particular stimulus actually triggers a change in animal behavior. Estimation of behavioral states using the STEFTR method will allow us to identify the behavior-triggering stimulus by focusing on the timing and/or place of the behavioral transition. Similarly, large-scale recording of neural activities from moving animals in the laboratory itself is difficult to interpret. However, state estimation and feature extraction of the behavior will greatly facilitate the identification of neural activities that are associated with behavioral transitions and/or specific behavioral features.

## ACKNOWLEDGEMENTS

We thank Drs. André Brown, Katsuyoshi Matsushita, Ken-ichi Hironaka, Takuma Degawa, Daisuke Yamamoto, Soh Kohatsu and the Kimura laboratory members for suggestions and comments on this work. Nematode strains were provided by the Caenorhabditis Genetics Center (funded by the NIH Office of Research Infrastructure Programs P40 OD010440) and by National Bioresource Project funded by the Ministry of Education, Culture, Sports, Science and Technology (MEXT), Japan. This work was supported by Interdisciplinary graduate school program for systematic understanding of health and disease (for S.J.Y.), by the Tohoku Ecosystem-Associated Marine Science (for K. S.), by KAKENHI 25249020 and JP16H06536 (for K.H.), 24681006, JP16H01769, JP16H06541 (for K.Y.), 16H04655, 18H05069, and17K19450 (for A.K.), JP16H06542 (for S.H.), JP16H06539 (for T.M.), JP16H06545 (for K.D.K.) from the MEXT, by PRESTO JPMJPR14D8 (for S.H.) by JST, and by Osaka University Co-Creation Program (for T.M. and K.D.K.).

## AUTHOR CONTRIBUTIONS

S.J.Y, T.M. and K.D.K. designed the experiments, S.J.Y, K.O., K.I., N.K., T.K., D. T., Y.Y., Y.I., F.H., K.F., Y.T., A.Y.-U., K.H., K.S., K.Y., A.T., Y.I., A.K., S.H. and K.D.K. performed the experiments, S.J.Y. and T.M. analyzed the data, and S.J.Y. and K.D.K. wrote the manuscript. All authors reviewed the manuscript.

